# How we transmit memories to other brains: constructing shared neural representations via communication

**DOI:** 10.1101/081208

**Authors:** A. Zadbood, J. Chen, Y.C. Leong, K.A. Norman, U. Hasson

## Abstract

Humans are able to mentally construct an episode when listening to another person’s recollection, even though they themselves did not experience the events. However, it is unknown how strongly the neural patterns elicited by mental construction resemble those found in the brain of the individual who experienced the original events. Using fMRI and a verbal communication task, we traced how neural patterns associated with viewing specific scenes in a movie are encoded, recalled, and then transferred to a group of naïve listeners. By comparing neural patterns across the three conditions, we report, for the first time, that event-specific neural patterns observed in the default mode network are shared across the encoding, recall, and construction of the same real-life episode. This study uncovers the intimate correspondences between memory encoding and event construction, and highlights the essential role our common language plays in the process of transmitting one's memories to other brains.

## Introduction

Sharing memories of past experiences with each other is foundational for the construction of our social world. What steps comprise the encoding and sharing of a daily life experience, such as the plot of a movie we just watched, with others? To verbally communicate an episodic memory, the speaker has to recall and transmit via speech her memories of the events from the movie. At the same time, the listener must comprehend and construct the movie’s events in her mind, even though she did not watch the movie herself. To understand the neural processes that enable this seemingly effortless transaction, we need to study three stages: 1) the speaker’s encoding and retrieval [1,2]; 2) the linguistic communication from speaker to listener [3,4]; and 3) the listener’s mental construction of the events [5–7]. To date, there has been no work addressing the direct links between the processes of memory, verbal communication, and construction (in the listener’s mind) of a single real-life experience. Therefore, it remains a mystery how information from a past experience stored in one person’s memory is propagated to another person’s brain, and to what degree the listener’s neural construction of the experience from the speaker’s words resembles the original encoded experience.

To characterize this cycle of memory transmission, we compared neural patterns during encoding, spoken recall, and mental construction of each scene in a movie (Figure 1). To closely mimic a real-life scenario, the study consisted of ***movie-viewers*** who watched a continuous movie narrative, a person (***speaker***) watching and then freely verbally recalling the same movie, and finally naïve ***listeners***, who had never seen the movie, listened to the audio recording of the spoken description. We searched for scene-specific neural patterns common across the three conditions. To ensure the robustness of the results, the full study was replicated using a second movie. This design allowed us to map the neural processes by which information is transmitted across brains in a real-life context, and to examine relationships between neural patterns underlying encoding, communication, and construction.

**Fig 1:**
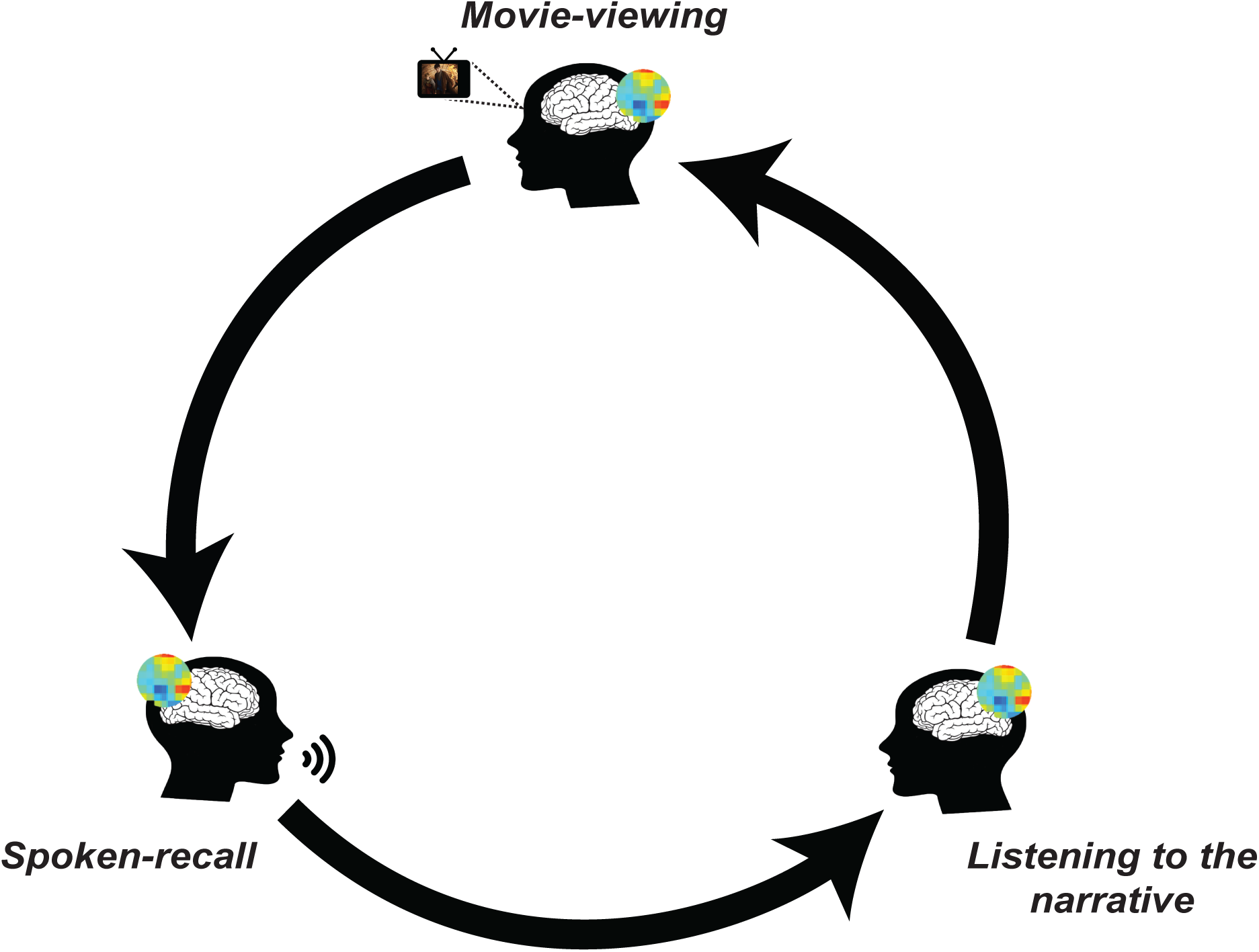
Circle of communication. Depiction of the entire procedure during sharing of an experience. Participants encode the movie and then reinstate it during recall. By listening to the audio recall, listeners construct the movie events in their mind. Mental representations related to the movie are shared throughout this cycle and transmitted across the brains via communication.

Why should we expect scene-specific neural patterns in high-order areas to be similar during the encoding, spoken recall, and mental construction of a given event? Resemblance between neural patterns elicited during encoding and retrieval have been shown in numerous studies using different types of stimuli [8–11, 2] over the past decade. More recently, it was demonstrated that ***scene-specific*** neural patterns elicited during encoding of complex natural stimuli (an audio-visual movie) are reinstated during free spoken recall [1]. Free spoken recall of a movie differs from the original experience (movie) in many aspects. Not only are they different modalities, but the content of each scene is altered during recall: recall is usually shorter and less detailed, with some elements emphasized and others minimized, based on the recaller’s preference or retrieval success. What are the neural processes underlying listening to such a recalled narrative? While no study has compared scene-specific patterns of brain responses during mental construction of a story with the patterns elicited during initial encoding or subsequent recall of the same event, recent studies suggest that the same areas that encode and retrieve episodic memories are also involved in the construction of imaginary and future events [12–20]. These areas include retrosplenial and posterior parietal cortices, ventromedial prefrontal cortex, bilateral hippocampus, and parahippocampal gyrus, known as default mode network [21,22]. Why are the same brain areas active during episodic encoding, retrieval, and mental construction? One possibility is that the same brain areas are involved in encoding, retrieval, and construction, but these areas assume different activity states during each process; in this case, one would expect that neural representations present during encoding and retrieval of specific scenes would not match those present during mental construction of those scenes. Another possibility is that the same neural activity patterns underlie the encoding, retrieval, and construction of a given scene. This hypothesis has never been tested.

Our communication protocol (Figure 1) provides a testbed for this latter hypothesis. In our experiment, during the spoken recall phase, the speaker must retrieve and reinstate her episodic memory of the movie events. At the same time, the listeners, who never experienced the movie events, must construct (imagine) the same events in their minds. Thus, if the same neural processes underlie both retrieval and construction, then we predict that similar activity patterns will emerge in the speaker’s brain and the listener’s brains while recalling/constructing each event. Furthermore, if the speaker successfully communicates her experiences of the original events to the listeners, then we should predict similarity between the neural patterns during the encoding phase (movie viewing) and construction phase (listening to the verbal description without viewing).

In the current study we witness, for the first time, how an event-specific pattern of activity can be traced throughout the communication cycle: from encoding, to spoken recall, to comprehending and constructing (Figure 1). Our work reveals the intertwined nature of memory, mental construction, and communication in real-life settings, and explores the neural mechanisms underlying how we transmit information about real-life events to other brains.

## Results

Eighteen participants watched a 25-minute audiovisual movie (from the first episode of BBC’s ***Merlin***) while undergoing fMRI scanning (***movie-viewing***, Figure 2-A). One participant separately watched the movie and then recalled it aloud inside the scanner (unguided, without any experimenter cues) and her spoken description of the movie was recorded (***spoken-recall***). Another group of participants (N = 18) who were naïve to the content of the movie listened to the recorded narrative (***listening***). The entire procedure was repeated with a second movie (from the first episode of BBC’s ***Sherlock***), with the same participant serving as the speaker. This design allowed us to internally replicate each of our findings and demonstrate the robustness of our results.

**Fig 2:**
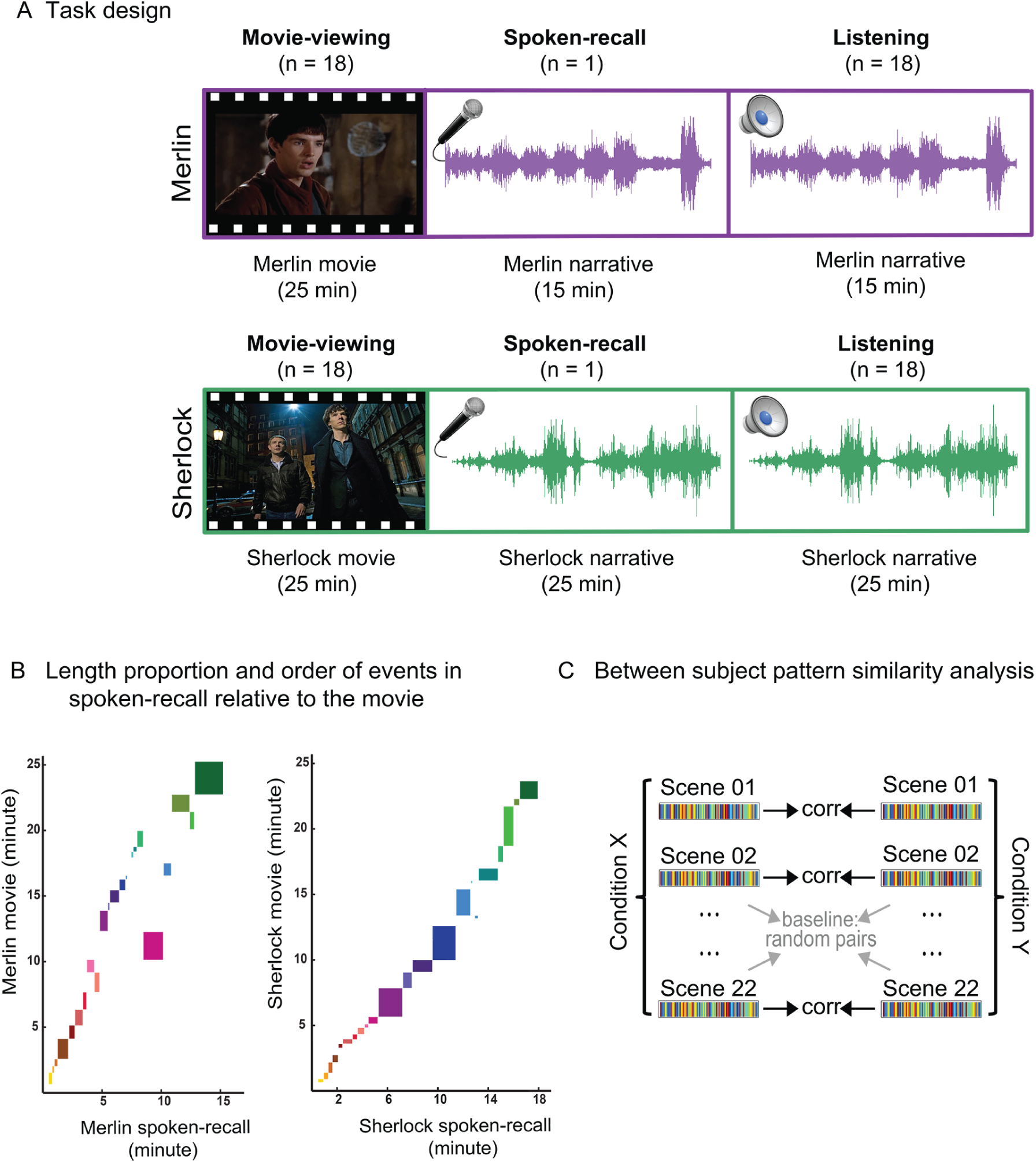
Experiment design and analysis A. 18 participants watched a 25-minute audiovisual movie (Merlin) while undergoing fMRI scanning (movie-viewing). One participant separately watched the movie and then recalled it inside the fMRI scanner and her spoken description of the movie was recorded (spoken-recall). Then a group of 18 participants who were naïve to the content of the movie listened to the recorded narrative. The entire procedure was repeated with a second movie (Sherlock) by recruiting new groups of participants. **B.** Depiction of the length of each event in the movie (y-axis) relative to its corresponding event (if remembered) in spoken-recall (x-axis) for each movie. Each box denotes a different event. Boxes that are out of the continuous diagonal string of events depict the events that were recalled in an order different from their original place in the movie. **C.** Schematic for the main analysis. Brain data were averaged within each scene in the data set of each condition (e.g. condition x = movie-viewing and condition Y = spoken-recall). Averaging resulted in a single pattern of brain response across the brain for each scene for each condition. Then these two patterns were compared and correlated using a searchlight method. Significant values were computed by shuffling the scene labels and comparing the non-matching scenes. Similar analyses were performed for all other comparisons (spoken-recall to listening, listening to movie-viewing)

## Pattern similarity between spoken-recall and movie-viewing

We first asked whether brain patterns elicited during ***spoken-recall*** (memory retrieval) were similar to those elicited during ***movie-viewing*** (encoding). To this end, we needed to compare corresponding content across the two datasets, i.e., compare brain activity as the ***movie-viewing*** participants encoded each movie event to the brain activity as the speaker recalled the same event during ***spoken-recall***. Previous work from our lab [1] has shown that neural patterns elicited by watching a movie are highly similar across participants at the individual scene level. Therefore, to increase the reliability of the movie-viewing-related patterns, we used the data from 18 viewers (not including the speaker’s viewing data) and compared them with the recall data in the speaker.

***Movie-viewing*** and ***spoken-recall*** data are not aligned across time-points; it took the speaker 15 minutes to describe the 25-minute Merlin movie, and 18 minutes to describe the 24-minute Sherlock movie (Figure 2-B). Therefore, data obtained during the watching of each movie (***movie-viewing***) were divided into 22 scenes (Figure 2-C), following major shifts in the narrative (e.g., location, topic, and/or time, as defined by an independent rater; see Methods for details). The same 22 scenes were identified in the audio recordings of the recall session based on the speaker’s verbal narration. Averaging time points within each scene provided a single pattern of brain response for each scene during recall. Pattern similarity analysis was conducted by calculating the Pearson correlation between the patterns elicited during movie-viewing and the patterns observed during the recall in a searchlight analysis (15 × 15 × 15 mm cubes centered on every voxel in the brain, [23,24]). This analysis reveals regions containing scene-specific reinstatement patterns, as statistical significance is only reached if matching scenes (same scene in movie and recall) can be differentiated from non-matching scenes [24]. In each voxel, scene labels were shuffled 10000 times and correlation was calculated which resulted in a null distribution. P-values were then calculated using this null distribution and were corrected for multiple comparisons using FDR (q < 0.05, two-tailed; see ***Methods***)

A large set of brain regions exhibited significant scene-specific similarity between the patterns of brain response during ***movie-viewing*** and ***spoken-recall***. Figure 3A shows the scene-specific ***movie-viewing*** to ***spoken-recall*** pattern similarity for the Merlin movie; Figure 3B replicates the results for the Sherlock movie. These areas included posterior medial cortex, medial prefrontal cortex, parahippocampal cortex, and posterior parietal cortex; collectively, these areas strongly overlap with default mode network (DMN). In the posterior cingulate cortex (PCC), a major region of interest (ROI) in the DMN, we observed a positive reinstatement effect in 17 of the 18 subjects in the ***Merlin*** condition (Fig. 3-C), and 18 out of the 18 subjects in the ***Sherlock*** condition (Fig. 3-D). The DMN has been previously shown to be active in episodic retrieval tasks [18,19,25]. Our finding of similar brain activity patterns between encoding and recall of a continuous movie narrative supports previous studies showing reinstatement of neural patterns during recall using simpler stimuli such as words, images, and short videos [2,8,11,26]. In addition, the result replicates a previous study from our lab that used a different dataset where both movie-viewing and recall were scanned for each participant [1].

**Fig 3:**
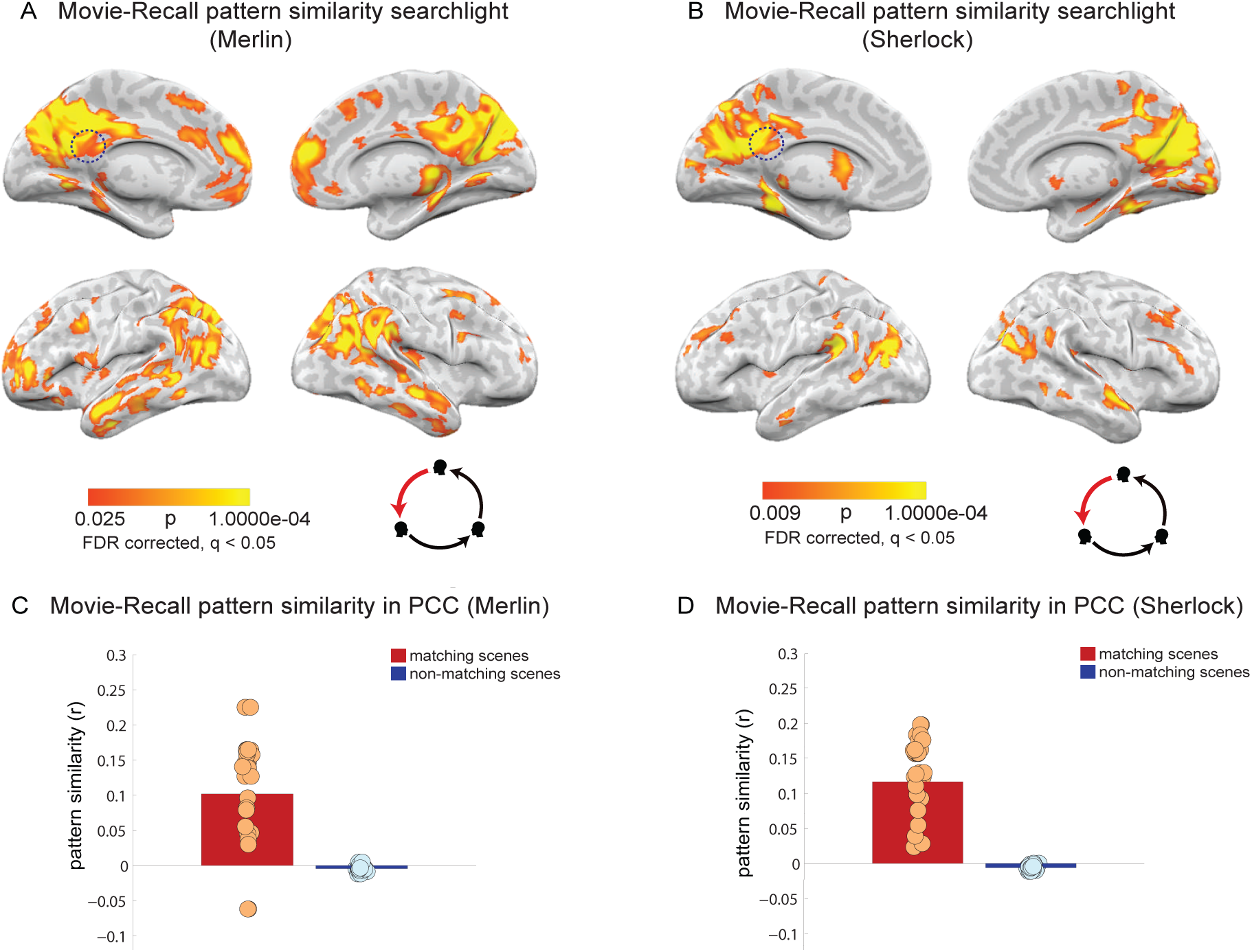
Movie-viewing to spoken-recall pattern similarity analysis A-B. Pattern similarity searchlight map, showing regions with significant between-participant, scene-specific correlations (p-values) between spoken-recall and movie-viewing (searchlight was a 5×5×5 voxel cube). Panel A depicts data for the Merlin movie and panel B depicts data for the Sherlock movie. Dotted circle shows the approximate location of the PCC ROI that was used in the analysis in panel C-D **C-D.** Pattern similarity (r-values) of each participant’s encoding (movie-viewing) data to the brain response during spoken-recall (in the speaker) in posterior cingulate cortex. Red bar shows average correlation of matching scenes and blue bar depicts average correlation of non-matching scenes, averaged across subjects. Circles depict values for individual subjects. Panel C depicts data for the Merlin movie and panel D depicts data for the Sherlock movie.

The above result shows that scene-specific brain patterns presented during the encoding of the movie were reinstated during the spoken free recall of the movie. Next we asked whether listening to a recording of the recalled (verbally described) movie would elicit these same event-specific patterns in an independent group of listeners who had never watched it (***listeners***).

## Pattern similarity between spoken-recall and listening

Previous studies have provided initial evidence for neural alignment (correlated responses in the temporal domain using inter-subject correlation) between the responses observed in the speaker’s brain during the production of a story and the responses observed in the listener’s brain during the comprehension of the story [3,4]. Moreover, it has been shown that higher speaker-listener neural coupling predicts successful communication and narrative understanding [3]. However, it is not known whether similar scene-specific ***spatial*** patterns will be observed across communicating brains, and where in the brain such similarity exists. To test this question, we implemented the same method as explained in the previous section (also see ***Methods***); however, for this analysis we correlated the average scene-specific neural patterns observed in the speaker’s brain during spoken recall with the average scene-specific neural patterns observed in the listeners’ brains as they listened to a recording of the spoken recall. Previous work suggests that during communication, the neural responses observed in the listener follows the speaker’s neural response timecourses with a delay of a few seconds [3,4,27]. To see whether this response lag was also present in our listeners’ brains, we calculated the correlation in PCC between the scene-specific neural patterns during ***spoken-recall*** and ***listening*** in the spatial domain, with TR-by-R shifting of listeners’ neural timecourses. Figure S1-A depicts the r values in the PCC ROI as the TR shift in the listeners was varied from −20 to 20 TRs (−30 to 30 seconds). In agreement with prior findings, we observed a lag between ***spoken-recall*** and ***listening**.* In the Merlin movie correlation peaked (r = 0.17) at a lag of ∼5 TRs (7.5 seconds). A similar speaker-listener peak lag correlation at ∼5 TRs was replicated in the ***listeners*** of the Sherlock movie (Fig. S1-B). To account for the listeners’ lag response, we used this 5 TR lag across the entire brain in all analyses.

We observed significant scene-specific correlation between the speaker’s neural patterns during the spoken recall and the listeners’ neural patterns during speech comprehension. Scene-specific neural patterns were compared between the ***spoken-recall*** and ***listening*** conditions using a searchlight and were corrected for multiple comparisons using FDR (q<0.05). Figure 4A shows the scene-specific ***spoken-recall*** to ***listening*** pattern similarity for the Merlin movie; Figure 4B replicates the results for the Sherlock movie. Similarity was observed in many of the areas that exhibited the memory reinstatement effect (movie-spoken recall correlation, Figure 3), including angular gyrus, precuneus, retrosplenial cortex, posterior cingulate cortex and mPFC.

**Fig 4:**
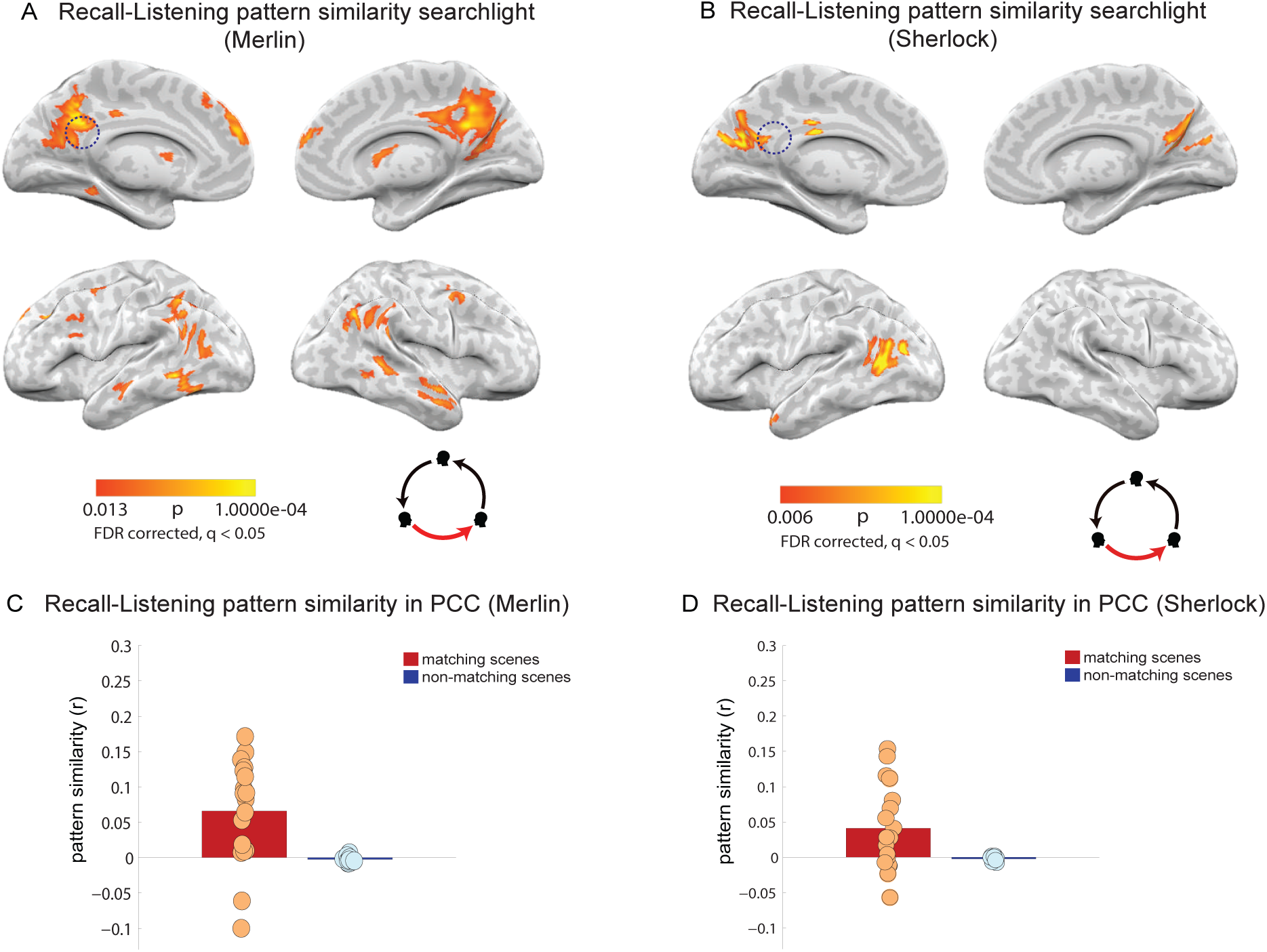
Spoken-recall to listening pattern similarity analysis A-B. Pattern similarity searchlight map, showing regions with significant between-participant, scene-specific correlations (p-values) between spoken-recall and listening (searchlight was a 5×5×5 voxel cube). Panel A depicts data for the Merlin movie and panel B depicts data for the Sherlock movie. Dotted circle shows the approximate location of the PCC ROI that was used in the analysis in panel CX-D **C-D.** Pattern similarity (r-values) of each participant’s listening data to the brain response during spoken-recall (in the speaker) in posterior medial cortex. Red bar shows average correlation of matching scenes and blue bar depicts average correlation of non-matching scenes, averaged across subjects. Circles depict values for individual subjects. Panel C depicts data for the Merlin movie and panel D depicts data for the Sherlock movie.

**Fig S1:**
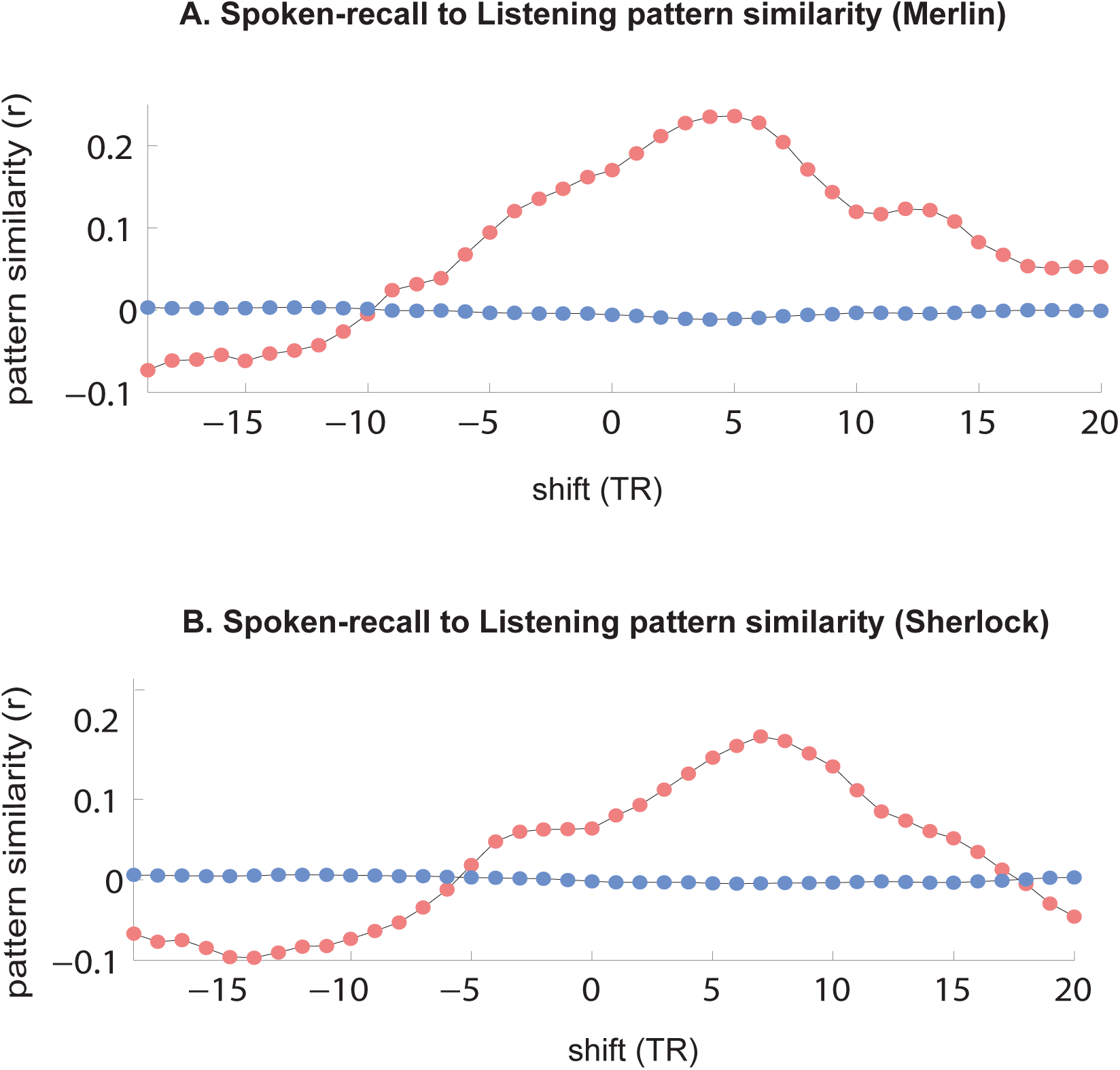
Pattern similarity in PCC with shifting of the listening data. This figure depicts the r values for pattern similarity between the spoken-recall data and average of all listeners’ listening data on y axis. Listening data has been shifted −20 to +20 TRs (x-axis) before calculating pattern similarity. R values for each shift are depicted as a separate dot. R values peak at TR = 5.

## Pattern similarity between listening and movie-viewing

So far we have demonstrated that event-specific neural patterns observed during encoding in high-order brain areas were reactivated in the speaker’s brain during spoken recall; and that some aspects of the neural patterns observed in the speaker were induced in the listeners’ brains while they listened to the spoken description of the movie. If speaker-listener neural alignment is a mechanism for transferring event-specific neural patterns encoded in the memory of the observer to the brains of naive listeners, then we predict that the neural patterns in the listeners’ brains during the construction of each event will resemble the movie-viewers’ neural patterns during each scene. To test this, we compared the patterns of brain responses when people listened to a verbal description of that event (***listening***) with those when people encoded the actual event while watching the movie (***movie-viewing***).

We found that the event-specific neural patterns observed as participants watched the movie were significantly correlated with neural patterns of naïve listeners who listened to the spoken description of the movie. Figure 5A shows the scene-specific ***listening*** to ***movie-viewing*** pattern similarity for the Merlin movie; Figure 5B replicates the results for the Sherlock movie. Similarity was observed in many of the same areas that exhibited memory reinstatement effects (***movie-viewing*** to ***spoken-recall*** correlation Figures 3) and speaker-listener alignment (Figures 4), including angular gyrus, precuneus, retrosplenial cortex, posterior cingulate cortex and mPFC. Computing the scene-specific ***listening*** to ***movie-viewing*** pattern similarity within the same PCC ROI shows that effect was positive for each of the individual subjects in each of the movies (Figure 5C-D).

**Fig 5:**
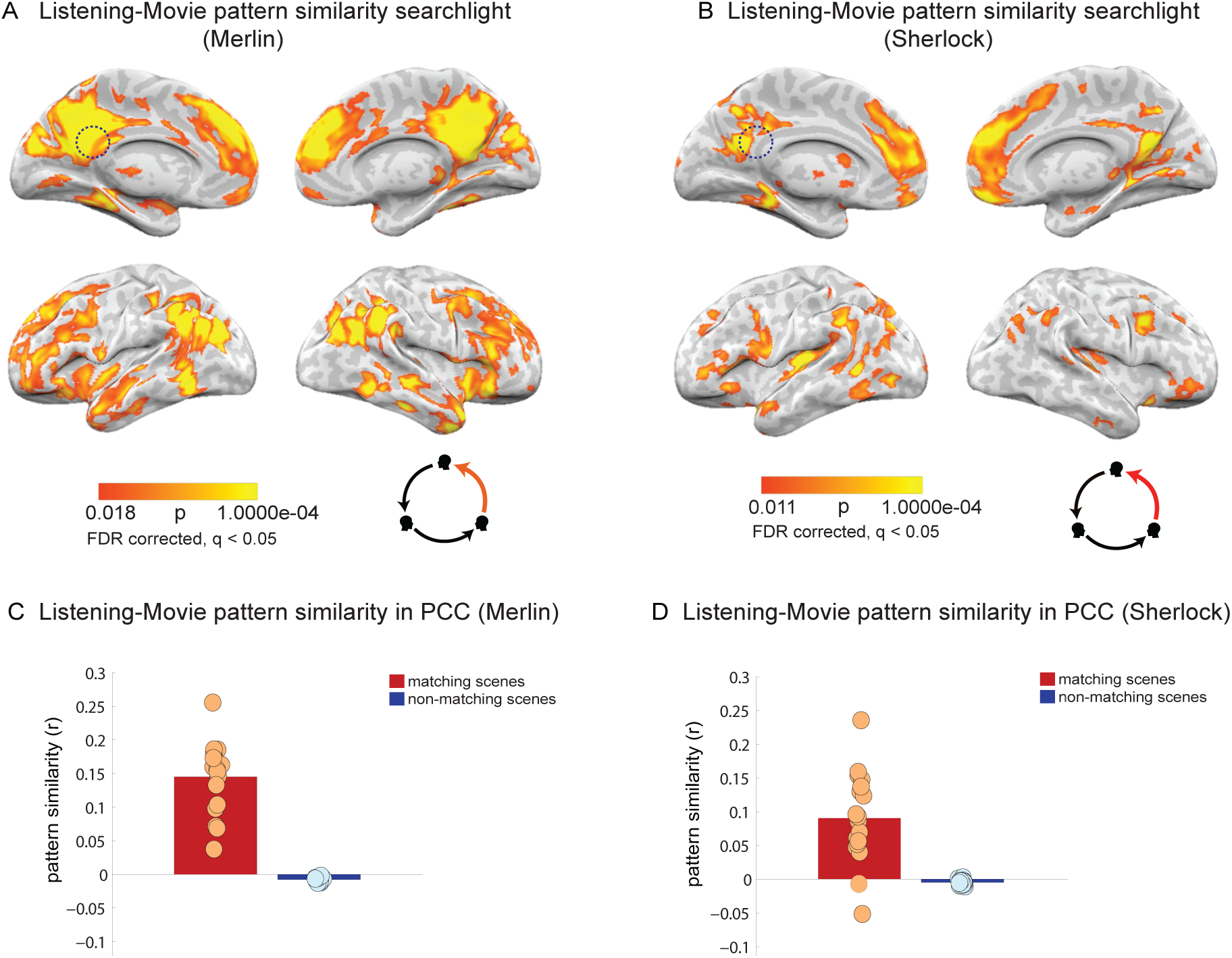
Listening to movie-viewing pattern similarity analysis A-B. Pattern similarity searchlight map, showing regions with significant between-participant, scene-specific correlations (p-values) between movie-viewing and listening (searchlight was a 5×5×5 voxel cube). Panel A depicts data for the Merlin movie and panel B depicts data for the Sherlock movie. Dotted circle shows the approximate location of the PCC ROI that was used in the analysis in panel C-D **C-D.** Pattern similarity (r-values) of each participant’s movie-viewing data to the average of all other listeners in posterior medial cortex. Red bar shows average correlation of matching scenes and blue bar depicts average correlation of non-matching scenes, averaged across subjects. Circles depict values for individual subjects. Panel C depicts data for the Merlin movie and panel D depicts data for the Sherlock movie.

**Fig S2:**
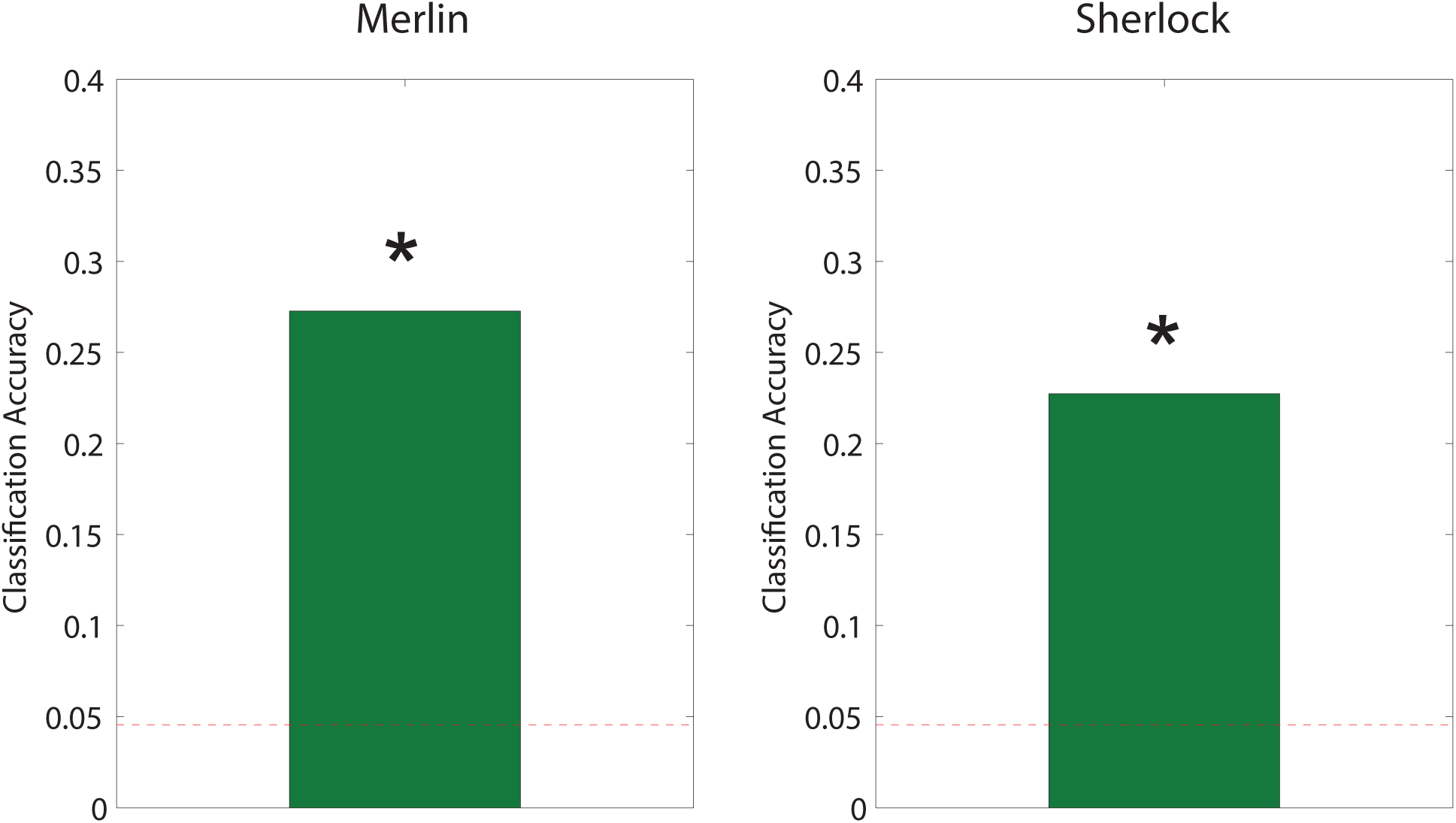
Viewing to Listening Scene Classification. Bar graphs depict overall classification accuracy between viewing and listening. This shows that, on average, a given scene could be correctly distinguished from other scenes even across modalities. The red line indicates the probability of correctly identifying a scene among 22 scenes based on guessing alone (4.5%). Based on a permutation test, the average classification accuracy for both movies was well above chance (p=0.0002 and p=0.001 for Merlin and Sherlock respectively).

To confirm that the relationship between the viewing and listening patterns was scene-specific, we assessed whether we could classify ***which scene*** participants were hearing about (in the listening condition) by matching scene-specific patterns from the listening condition to scene-specific patterns from the viewing condition. We created average patterns in posterior cingulate cortex for each scene separately for viewing and listening groups. On average, the neural pattern observed during movie-viewing of a particular scene was most similar to the pattern observed when listening to a verbal description of the scene (average classification accuracy for Merlin = 27%, p = 0.0002 1-tailed, Sherlock = 22%, p = 0.001 1-tailed, chance level = 4.5%, Fig S-2), even though participants listening to the verbal description had not previously seen the movie

## Relationship between pattern similarity and behavioral performance

Given that the speaker’s success in transmitting her memories may vary across listeners, we next asked whether the level of correlation between the neural responses of each listener and the speaker’s neural responses while encoding the movie can predict the listeners’ comprehension level. To test this question, we looked at the posterior cingulate cortex (PCC). The PCC was chosen as the region of interest since previous research has shown that the strength of similarity between spatial patterns of brain response during encoding and rehearsal in this area could predict the subsequent memory performance [2]. Indeed, within the posterior cingulate cortex, speaker-listener neural alignment (correlation) predicted the level of comprehension in the listeners, as measured with an independent post-scan test of memory and comprehension (Figure 6-A and 6-C, R = 0.71 and P = 0.001 for the Merlin movie, R = 0.54 and P = 0.022 for the Sherlock movie).

**Fig 6:**
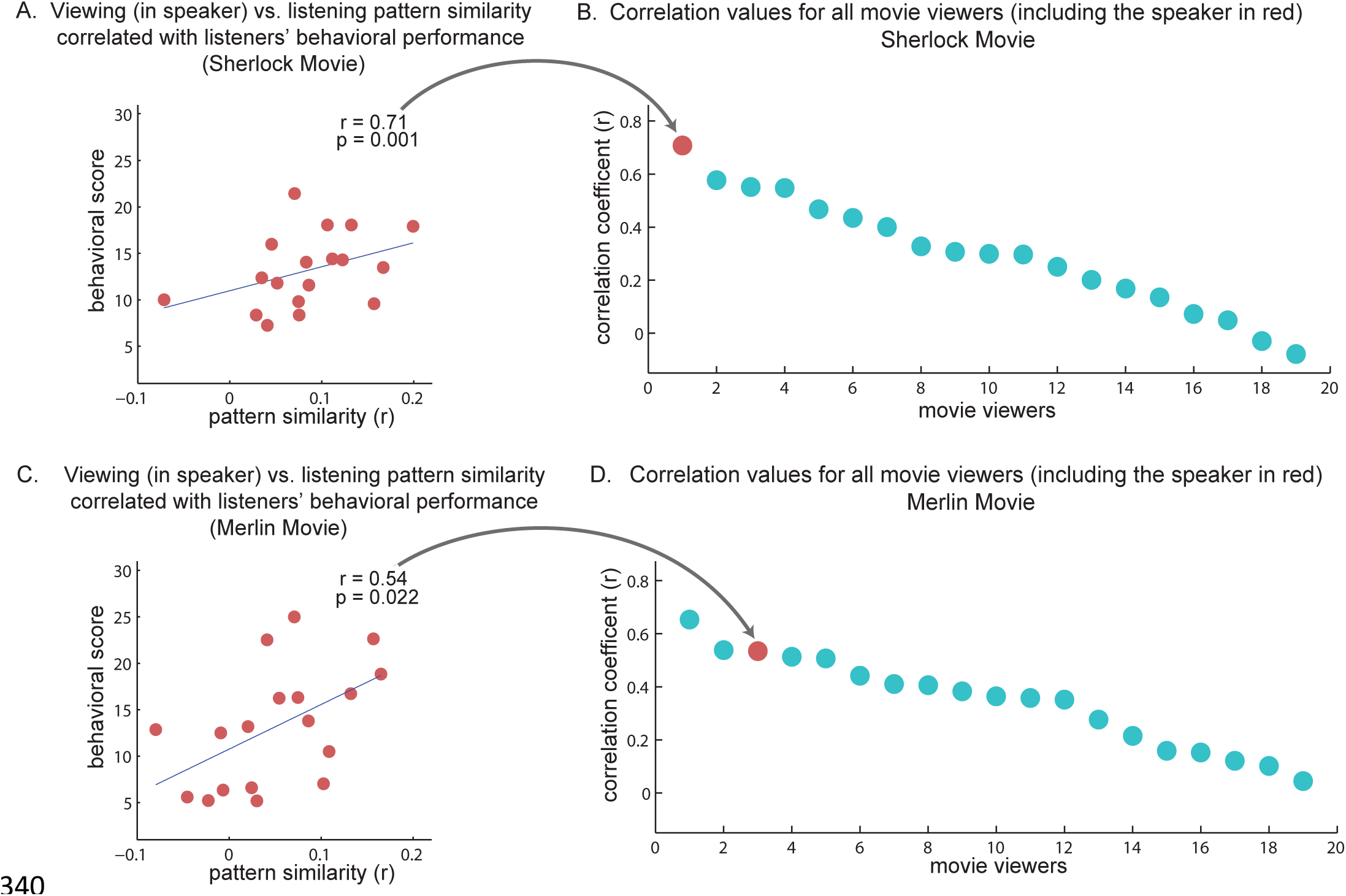
Pattern similarity of movie viewers to listeners – relationship to comprehension level. A,C. Correlation between the comprehension score of listeners and degree of similarity between the speaker neural responses during the encoding phase (i.e. while watching the movie) and all listeners in PCC, for each movie. **B,D** Rank order correlation values of the same analysis as in A,C for each of the viewers (including the speaker, red circle). Note that correlating the listeners’ brain responses with the actual speaker’s brain responses during encoding phase better predicted comprehension levels than the correlation with other viewers.

Different people could vary in the way they encode and memorize the same events in the movie. These idiosyncrasies would then be transmitted to listeners when a particular speaker recounts her memory. A successful transmission of a particular episodic memory, therefore, may entail a stronger correspondence between the neural responses of the listeners with those of the ***speaker watching the movie***, as opposed to with other viewers watching the same movie. To test this hypothesis, we compared the listeners’ comprehension levels with the correlation between neural patterns in posterior cingulate cortex of each movie viewer (including the speaker, N = 19) and each listener during listening. We observed that the listeners’ comprehension levels were predicted the best when we compared the listeners’ neural patterns with those of the ***actual speaker viewing the movie***, relative to all other 18 viewers (Figure 6-A); and among the top three in the replication study (Figure 6-C). These results indicate that during successful communication the neural responses in the listeners’ brains were aligned with neural responses observed in the speaker’s brain during encoding (viewing) the movie, even before recall had begun.

## Shared neural response across three conditions (triple shared pattern analysis)

In Figures 3, 4 and 6 we show the pairwise correlations between encoding, speaking, and constructing. The areas revealed in these maps are confined to high order areas, which overlap with the default mode network, and include the TPJ, angular gyrus, retrosplenial, precuneus, posterior cingulate cortex and mPFC. Such overlap suggests that there are similarities in the neural patterns, which are shared at least partially, across conditions. Correlation, however, is not transitive (beside the special case when the correlation values are close to 1). That is, if x is correlated with y, y is correlated with z, and z is correlated with x, one can’t conclude that a shared neural pattern is common across all three conditions. To directly quantify the degree to which neural patterns are shared across the three conditions, we developed a new, stringent three-way similarity analysis to identify shared event-specific neural patterns across all three conditions (movie encoding, spoken recall, naïve listening). The analysis looks for shared neural patterns across all conditions, by searching for voxels that fluctuate together (either going up together or down together) in all three conditions (see methods for details). Figure 7A shows all areas in which the scene-specific neural patterns are shared across all three conditions in the Merlin movie;; Figure 7B replicates the results in the Sherlock movie. These areas substantially overlap with the pairwise maps (Figs 3, 4 and 6), thereby indicating that similarities captured by our pairwise correlations include patterns that are shared across all three conditions. Note that the existence of shared neural patterns across conditions does not preclude the existence of additional response patterns that are shared across only two of the three conditions (e.g. shared responses across the speaker-listener which are not apparent during movie encoding), and revealed in the pair-wise comparisons (Figures 3, 4 and 6).

**Fig 7:**
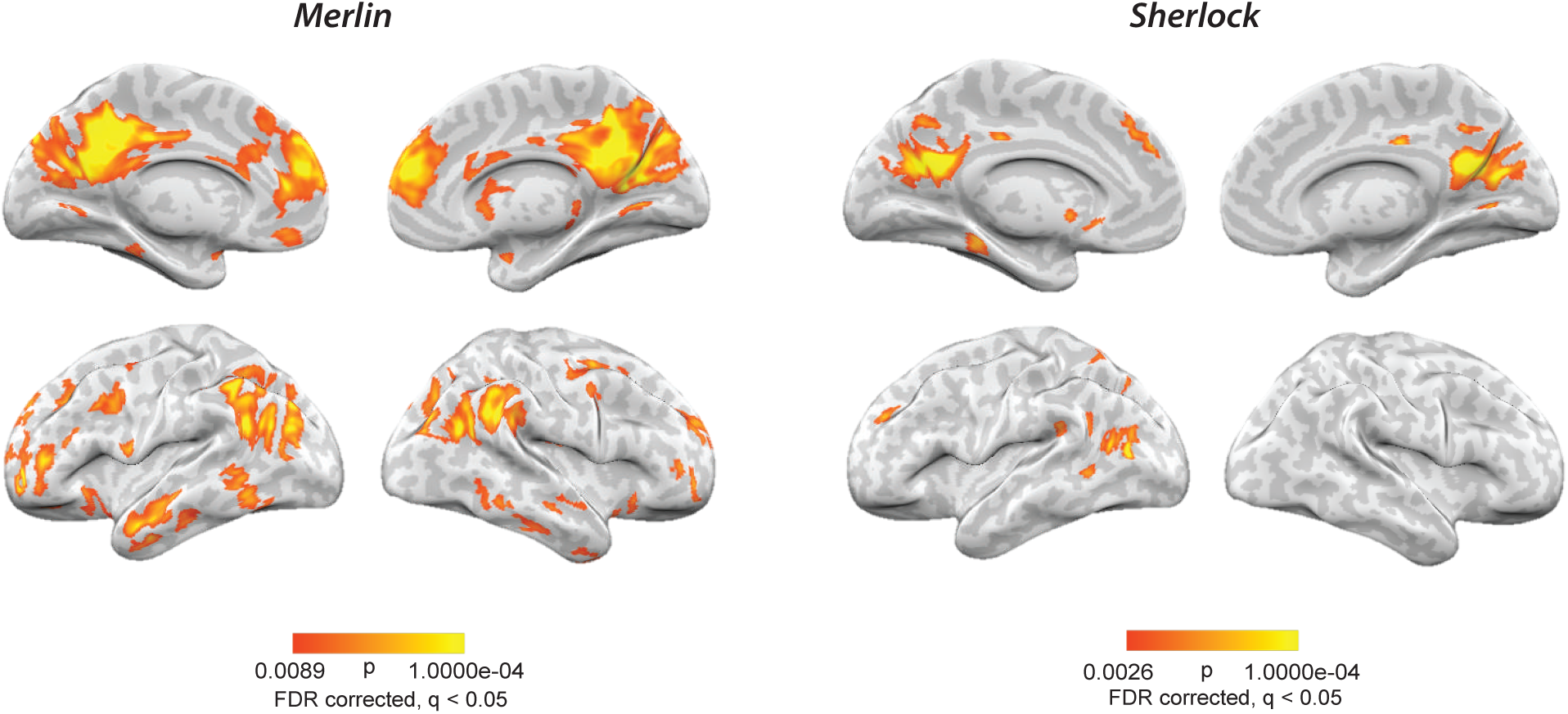
Shared neural patterns across all conditions. Regions showing scene-specific pattern correlations across movie-viewing, spoken-recall, and listening for **A.** the Merlin movie **B.** the Sherlock movie.

## Discussion

This study reports, for the first time, that shared event-specific neural patterns are observed in the default mode network (DMN) during the encoding, reinstatement (spoken recall), and new construction of the same real-life episode. Furthermore, across participants, higher levels of similarity between the speaker’s neural patterns during movie viewing and the listeners’ neural patterns during mental construction were associated with higher comprehension of the described events in the listeners (i.e., successful “memory transmission”). Prior studies have shown that neural patterns observed during the encoding of a memory are later reinstated during recall [1,2,8–11]. Furthermore, it has been reported that the same areas that are active during recall are also active during prospective thinking and mental construction of imaginary events [14–17,28,29]. Our study is the first to directly compare scene-specific neural patterns observed during ***mental construction (imagination) of a verbally-described but never experienced event*** directly to patterns elicited during ***audio-visual perception of the original event***. This comparison, which was necessarily performed across-participants, revealed brain areas throughout the DMN, including posterior medial cortex, mPFC, and angular gyrus, where spatial patterns were shared across both spoken recall and mental construction of the same event.

Why do we see such a strong link between memory encoding, spoken recall and construction? By identifying these shared event-specific neural patterns, we hope to illustrate an important purpose of communication: to transmit and share one’s thoughts and experiences with other brains. In order to transmit memories to another person, a speaker needs to convert between modalities, using speech to convey what she saw, heard, felt, smelled, or tasted. In our experimental setup, during the spoken recall, the speaker focused primarily on the episodic narrative (e.g., the plot, locations and settings, character actions and goals), rather than on fine sensory (visual and auditory) details. Accordingly, ***movie-viewing*** to ***spoken-recall*** pattern correlations were not found in low level sensory areas, but instead were located in high level DMN areas, which have been previously found to encode amodal abstract information [30–32]. Future studies could explore whether the same speech-driven recall mechanisms can be used to reinstate and transmit detailed sensory memories in early auditory and visual cortices.

Spoken words not only enabled the reinstatement of scene-specific patterns during recall, but also enabled the construction of the same events and neural patterns as the listeners imagined those scenes. For example, when the speaker says “Sherlock looks out the window, sees a police car, and says, well now it’s four murders”, she uses just a few words to evoke a fairly complex situation model. Remarkably, a few brief sentences such as this are sufficient to elicit neural patterns, specific to this particular scene, in the listener’s DMN that significantly resemble those observed in the speaker’s brain during the scene encoding. Thus, the use of spoken recall in our study exposes the strong correspondence between memories (event reconstruction) and event construction (imagination). This intimate connection between memory and imagination [13,28,33,34] allows us not only to share our memories with others, but also to invent and share imaginary events with others. Areas within the DMN have been proposed to be involved in creating and applying “situation models” [35,36], and changes in the neural patterns in these regions seem to mark transitions between events or situations [37,38]. An interesting possibility is that the (re)constructed “situation model” is the “unit” of information transferred from the speaker to the listener, a transfer made compact and efficient by taking advantage of their shared knowledge.

We showed that, despite the differences between the verbal recall and the movie, listening to a recalled narrative of the movie triggered mental construction of the events in the listeners’ brain and enabled them to partly experience a movie they have never watched. Similarity between patterns of brain response during perception and imagination has been reported before [39–42]. These studies have mostly focused on visual or auditory imagery of static objects, scenes and sounds. In our study, we directly compared, for the first time, scene-specific neural patterns during mental construction of rich episodic content, which describes the actions and intentions of characters as embedded in real-life dynamical movie narrative.

In agreement with the hypothesis that the speaker’s verbal recall transmitted her own idiosyncratic memory of the movie, we found the listeners correlated better with the speaker’s neural patterns during the encoding of the movie, relative to neural responses in other viewers that watched the movie. Furthermore, the ability of the speaker to successfully transmit her memories can vary as a function of how successful the listeners are in constructing the information in their minds. And indeed we observed that the strength of speaker-listener neural alignment correlated with the listeners’ comprehension as measured by a post-scan memory and comprehension tests. Taken together, these results suggest that the alignment of brain patterns between the speaker and listeners can capture the quality of transfer of episodic memories across brains. This finding extends previous research that showed a positive correlation between communication success and speaker-listener neural coupling in the temporal domain [3,4,27] in posterior medial cortex, and is also consistent with research showing that higher levels of encoding-to-recall pattern similarity in posterior cingulate cortex positively correlate with behavioral memory measures [2]. Our result highlights the importance of the subjectivity and uniqueness of the original experience (how each person perceives the world, which later on affects how they retrieve that information) in transmission of information across different brains.

What causes some listeners to have weaker or stronger correlation with the speaker’s neural activity? Listeners may differ in terms of their ability to construct and understand second hand information that is transmitted by the speaker. The speaker’s recall is biased toward those parts of the movie which are more congruent with her own prior knowledge, and the listener's comprehension and memory of the speaker’s description is also influenced by his/her own prior knowledge [34,43–45]. Thus, the coupling between speaker and listener is only possible if the interlocutors have developed a shared understanding about the meaning and proper use of each spoken (or written) sign [46–48]. For example, if instead of using the word “police officers” the speaker uses the British synonym “bobbies”, she is likely to be misaligned with many of the listeners. Thus, the construction of the episode in the listeners’ imagination can be aligned with speaker’s neural patterns (associated with the reconstruction of the episode) only if both speaker and listener share the rudimentary conceptual elements that are used to compose the scene.

Finally, it is important to note that information may change in a meaningful or useful way as it passes through the communication cycle; the three neural patterns associated with encoding, spoken recall, and construction are similar but not identical. For example, in a prior study we documented systematic transformations of neural representations between movie encoding and movie recall [1]. In the current study, we observed that the verbal description of each scene seemed to be compressed and abstracted relative to the rich audio-visual presentation of these events in the movie. Indeed, at the behavioral level, we found that most of the scene recalls were shorter than the original movie scene (e.g., in our study it took the speaker ∼15-18 minutes to describe a ∼25-minute movie). Nevertheless, the spoken descriptions were sufficiently detailed to elicit replay of the sequence of scene-specific neural patterns in the listeners’ DMNs. Because the DMN integrates information from multiple pathways [49,50], we propose that, as stimulus information travels up the cortical hierarchy of timescales during encoding, from low-level sensory areas up to high-level areas, a form of compression takes place [51]. These compressed representations in the DMN are later reactivated (and perhaps further compressed) using spoken words during recall. It is interesting to note that the listeners may benefit from the speaker’s concise speech, as it allows them to bypass the step of actually watching the movie themselves. This may be an efficient way to spread knowledge through a social group (with the obvious risk of missing on important details), as only one person needs to expend the time and run the risks in order to learn something about the world, and can then pass that information on to others.

Overall, this study tracks, for the first time, how real-life episodes are encoded and transmitted to other brains through the cycle of communication. Sharing information across brains is a challenge that the human race has mastered and exploited. This study uncovers the intimate correspondences between memory encoding and narrative construction, and highlights the essential role that our shared language plays in that process. By demonstrating how we transmit mental representations of previous episodes to others through communication, this study lays the groundwork for future research on the interaction between memory, communication, and imagination in a natural setting.

## Materials and Methods

### Stimuli

We used two audio-visual movies, the first episodes of Sherlock BBC (24-min length) and Merlin (25-min length) BBC. These movies were chosen to have similar levels of action, dialogue, and production quality. Audio recordings were obtained from a participant who watched and recounted the two movies in the scanner (free-recall). The outcome was an 18-min audio recording of the Sherlock story, and a 15-min audio recording of the Merlin story. Thus the stimuli consisted of a total of two movies (Sherlock and Merlin) and two corresponding audio recordings. This allowed us to internally replicate the results across the two datasets.

### Subjects

A total of 52 participants (age 18 – 45) who were all right-handed native English speakers with normal or corrected to normal vision were scanned. Before contacting participants, their previous exposure to both movie stimuli was screened and only people without any self-reported history of watching either of the two movie stimuli were recruited. From the total group, 4 were dropped due to head motions larger than 3 mm (voxel size), 1 was dropped due to anomalous anatomy, 4 fell asleep, 5 were dropped due to failure in post scan memory test (recall levels < 1.5 SD below the mean), and 2 were dropped who had watched the movie but did not report it before the scan session. Subjects who were dropped due to poor recall had scores close to zero (Merlin scores: max = 25, min = 0.4, mean = 11.9, std = 7.1 Sherlock scores: max = 21.4, min = 0, mean = 11.18, std = 5.6). We acquired informed consent from all participants, which was approved by Princeton University Institutional Review Board.

### Procedure

***Experimental design***. One participant watched both movies (Sherlock and Merlin) in the scanner in separate sessions and recalled them out loud while being scanned. She was instructed before the scan that she would be asked to recall the movies afterward. There were two main runs in the experiment. During the first run, participants watched either the Sherlock or Merlin movie (***movie-viewing***). During the second run, participants listened to an audio description of the movie they had not watched (***listening***). After the main experiment, participants listened to a short audio stimulus (15 minutes) in the scanner. Data from this run were collected for a separate experiment and was not used in this paper. Participants were randomly assigned to watch Sherlock (n = 18) or Merlin (n =18). Sound level was adjusted separately for each participant to assure a complete and comfortable understanding of the stimuli. An anatomical scan was performed at the end of the scan session. Before the experiment, participants were instructed to watch and/or listen to the stimuli carefully and were told that there would be memory tests for each part separately.

There was no memory task (or any task) inside the scanner and there was no specific instruction about fixating to the center. Participants were asked to watch the stimuli through the mirror which was reflecting the rear screen. The movie was projected to this screen located at the back of the magnet bore via a LCD projector. In-ear headphones were used for the audio stimuli. Eye-tracking was performed during all the runs (recording during the movie, observing the eye during the audio) using iView X MRI-LR system (SMI Sensomotoric Instruments). Eye-tracking was implemented to ensure that participants were paying full attention and not falling asleep. They were asked to keep their eyes open even during the audio runs (no visual stimuli). The movie and audio stimuli were presented using Psychophysics Toolbox [http://psychtoolbox.org] in MATLAB, which enabled us to coordinate the onset of the stimuli (movie and audio) and data acquisition.

***MRI acquisition:*** MRI data was collected on a 3T full-body scanner (Siemens Skyra) with a 16-channel head coil. Functional images were acquired using a T2^*^-weighted echo planar imaging (EPI) pulse sequence (TR 1500 ms, TE 28 ms, flip angle 64, whole-brain coverage 27 slices of 4 mm thickness, in-plane resolution 3 × 3 mm2, FOV 192 × 192 mm2). Anatomical images were acquired using a T1-weighted magnetization-prepared rapid-acquisition gradient echo (MPRAGE) pulse sequence (0.89 mm3 resolution). Anatomical images were acquired in an 8-minute scan after the functional scan with no stimulus on the screen.

### Post-scan behavioral memory test

Memory performance was evaluated using a free recall test in which participants were asked to write down the events they remembered from the movie and audio recording with as much detail as possible. There was no time limitation and they were asked to ensure they wrote everything that they remembered. Three independent raters were asked to read the transcripts of participants’ free recalls and to assign memory scores to each participant. The raters were given general instructions to assess the quality of the comprehension and accuracy of each response, and a few examples. They reported a score for each participant and these numbers were normalized to the same scale across the three raters. Ratings generated by of the three raters were highly correlated (Cronbach’s Alpha = 0.85 and 0.87 for Merlin and Sherlock respectively) and averaged to be used in further analysis.

### Data analysis

Preprocessing was performed in FSL [http://fsl.fmrib.ox.ac.uk/fsl], including slice time correction, motion correction, linear detrending, and high-pass filtering (140 s cutoff). These were followed by coregistration and transformation of the functional volumes to a template brain (MNI). The rest of the analysis was coded and performed using Matlab software (MathWorks). All the time courses were despiked before further analysis. We briefly review the analytical methods and objectives. Before running the searchlight analysis, brain time-courses were averaged within each scene for all the participants and conditions.

#### Pattern similarity searchlight

For each searchlight analysis [23], pattern similarity was computed in 5 × 5 × 5 voxel cubes (15 × 15 × 15 mm) by placing the center of the cube on every voxel across the brain and calculating the correlation between patterns. Significance thresholds were calculated using a permutation method [24] by shuffling the scene labels and correlating non-matched scenes to create a null distribution of r-values; the p-value was extracted from this distribution (2-tailed). This procedure was implemented for all the searchlight cubes for which 50% or more of their volume was inside the brain. Thus individual p values were generated for each voxel (center of searchlight cube) and were corrected for multiple comparisons using False Discovery Rate [52], q < 0.05. This analysis aims to confirm the event-specificity of our findings by demonstrating that correlation between matching scenes is significantly higher that non-matching scenes.

Encoding to recall pattern similarity was calculated by executing the searchlight analysis to compare the ***spoken-recall*** data with each subjects’ ***movie-viewing*** (encoding data) and then averaging across subjects. Pattern similarity analysis was performed across subjects. Therefore, the speaker’s movie viewing data was not included in the movie-viewing set. After performing the shuffling and permutation test, the average map was plotted with specific p-values for each voxel, with the threshold corrected using FDR (Figure 3.A-B). To compare ***speaking-to-listening***, the pattern similarity searchlight was used to compare the ***speaker***’s recall data with each of the ***listeners’*** listening data and then averaged and statistically thresholded (Figure 4.A-B). In ***listening-to-viewing*** condition, each ***viewer***’s data was correlated with the average of all the *listeners* listening data. The procedure was done for all the participants in the group and then statistical analysis and averaging was performed to compute the p-value maps (Figure 6.A-B). After averaging, maps were thresholded based on significance (FDR correction, q<0.05). For encoding to recall comparison, searchlight was restricted to voxels that exhibited reliable response to movie stimuli. This reliability was measured by inter subject correlation [53] of at least r = 0.1 (∼%70 of voxels in the brain). Recall to listening searchlight was restricted to voxels with ISC of at least r = 0.1 during listening (∼%20 of all brain voxels). In listening to movie viewing comparison we included voxels that were reliable (ISC at least 0.1) during either listening or movie viewing (∼%70 of voxels in the brain). Performing the searchlight without any voxel restriction resulted in similar results. FDR correction of unmasked maps did not change the p-values notably.

#### ROI-based pattern similarity

In addition, pattern similarity was separately calculated at subject level in posterior medial cortex ROI. This analysis was performed by calculating Pearson correlation between patterns of brain response across the entire ROI in each ***viewer*** to the ***speaker*** (Figure 3.C-D), each ***listener*** to the ***speaker*** (Figure 4.C-D), and each ***viewer*** to the average of all ***listeners*** (Figure 6.C-D). ROI level pattern similarity between the ***speaker*** and each ***listener*** was also computed in mPFC, and A1. Pattern similarity scores (correlation coefficients) for each ROI for each ***listener*** (from the ***speaker-listener*** correlation) were then correlated with that listener’s behavioral score (Figure 5). We used the posterior cingulate ROI from Shirer et al’s resting state connectivity atlas [54].

#### Behavioral correlation

To compute the behavioral correlation, pattern similarity in each viewer (including the speaker’s viewing) to each listener was calculated in PCC. These patterns similarity values for each viewer (18 values because of the correlation of each viewer with each of the 18 listeners) were then correlated with the listeners’ behavioral scores. Figure 5-B and Figure 5-C show the correlation for the speaker’s viewing and the listeners. Figure 5-A and Figure 5-C depict the sorted outcome r values of correlation for each movie viewer and show the speaker’s viewing in red as one of the highest values. To avoid the need to correct for multiple comparisons we did not test any other region. Furthermore, as with all other results in the paper, we run the same analysis on the second independent data set.

#### Classification analysis

To investigate the discriminability of neural patterns for individual scenes, we first averaged the time-course of brain response within each scene during movie viewing and listening. The patterns were then averaged across participants in each group to make an averaged pattern for viewing and an averaged pattern for listening. Pairwise correlation between the two groups for all 22 scenes was computed. Classification was considered successful if the pairwise correlation of any given scene between the movie and listening (matching scene) was higher than their correlation with any other scene (out of 22 possibilities, chance 4.5%). Overall accuracy was then computed by the number of scenes with the highest correlation (rank = 1 for matching scene) divided by the number of scenes. Chance was calculated by shuffling the scene labels and computing the pairwise correlation 10000 times. Null distribution was created using the classification accuracy of the shuffled scenes. Real classification accuracies of both movies stand beyond 99.9 percentile on this distribution.

#### Triple shared pattern searchlight

The triple shared pattern analysis was performed to directly compare the neural patterns across the three conditions (***movie-viewing, spoken-recall, listening***). We sought to find voxels within each searchlight cube that were correlated across the three conditions. For each scene, the brain response was z-scored across voxels (spatial patterns) within each cube. If the same patterns are present across conditions, then the z-scored activation value for a given voxel should have the same sign across conditions. To measure this property, we implemented the following computation: For a given voxel in each cube, if it showed all positive or all negative values across the three conditions, we calculated the product of the absolute values of brain response in that voxel. Otherwise (if a voxel did not exhibit all positive or negative signs across the three conditions), the product value was set to zero. The final value for each voxel was then created by averaging these product values across scenes. To perform significance testing, the order of scenes in each condition was randomly shuffled (separately for each condition) and then the same procedure was applied (calculating the product value and averaging). By repeating the shuffling 10000 times and creating the null distribution, p values were calculated for each voxel. The resulting p-values were then corrected for multiple comparisons using FDR (q < 0.05).

## Acknowledgments

We thank Christopher Baldassano for guidance on triple shared pattern analysis and his comments on the manuscript; We also thank Mor Regev, Yaara Yeshurun-Dishon and other members of the Hasson lab for scientific discussions, helpful comments and their support. This work was supported by The National Institutes of Health (R01-MH094480 and DP1 HD091948).

## References

1. Chen J, Leong YC, Honey CJ, Yong CH, Norman KA, Hasson U. Shared memories reveal shared structure in neural activity across individuals. Nat Neurosci. 2017;20: 115–125. doi: 10.1038/nn.4450.

2. Bird CM, Keidel JL, Ing LP, Horner AJ, Burgess N, Consolidation of Complex Events via Reinstatement in Posterior Cingulate Cortex. J Neurosci. 2015;35: 14426–14434. doi: 10.1523/JNEUROSCI.1774-15.2015.

3. Stephens GJ, Silbert LJ, Hasson U Speaker-listener neural coupling underlies successful communication. Proc Natl Acad Sci U S A. 2010; 107: 14425–14430. doi: 10.1073/pnas.1008662107.

4. Silbert LJ, Honey CJ, Simony E, Poeppel D, Hasson U Coupled neural systems underlie the production and comprehension of naturalistic narrative speech. Proc Natl Acad Sci. 2014; 111: E4687–E4696. doi: 10.1073/pnas.1323812111.

5. Chow HM, Mar RA, Xu Y, Liu S, Wagage S, Braun AR. Embodied comprehension of stories: interactions between language regions and modality-specific neural systems. J Cogn Neurosci. 2014;26: 279–295. doi: 10.1162/jocn_a_00487.

6. Mar RA. The neuropsychology of narrative: story comprehension, story production and their interrelation. Neuropsychologia. 2004;42: 1414–1434. doi: 10.1016/j.neuropsychologia.2003.12.016.

7. Kosslyn SM, Ganis G, Thompson WL. Neural foundations of imagery. Nat Rev Neurosci. 2001;2: 635–642. doi: 10.1038/35090055.

8. Polyn SM, Natu VS, Cohen JD, Norman KA. Category-specific cortical activity precedes retrieval during memory search. Science. 2005;310: 1963–1966. doi: 10.1126/science.1117645.

9. Johnson JD, McDuff SGR, Rugg MD, Norman KA. Recollection, familiarity, and cortical reinstatement: a multivoxel pattern analysis. Neuron. 2009;63: 697–708. doi: 10.1016/j.neuron.2009.08.011.

10. Buchsbaum BR, Lemire-Rodger S, Fang C, Abdi H, The Neural Basis of Vivid Memory Is Patterned on Perception. J Cogn Neurosci. 2012;24: 1867–1883. doi: 10.1162/jocn_a_00253.

11. St-Laurent M, Abdi H, Bondad A, Buchsbaum BR. Memory Reactivation in Healthy Aging: Evidence of Stimulus-Specific Dedifferentiation. J Neurosci. 2014;34: 4175–4186. doi: 10.1523/JNEUROSCI.3054-13.2014.

12. Hassabis D, Maguire EA. Deconstructing episodic memory with construction. Trends Cogn Sci. 2007;11: 299–306. doi: 10.1016/j.tics.2007.05.001.

13. Hassabis D, Maguire EA. The construction system of the brain. Philos Trans R Soc B Biol Sci. 2009;364: 1263–1271. doi: 10.1098/rstb.2008.0296.

14. Schacter DL, Addis DR, Buckner RL. Remembering the past to imagine the future: the prospective brain. Nat Rev Neurosci. 2007;8: 657–661. doi: 10.1038/nrn2213.

15. Addis DR, Wong AT, Schacter DL. Remembering the past and imagining the future: Common and distinct neural substrates during event construction and elaboration. Neuropsychologia. 2007;45: 1363–1377. doi: 10.1016/j.neuropsychologia.2006.10.016.

16. Spreng RN, Mar RA, Kim ASN. The common neural basis of autobiographical memory, prospection, navigation, theory of mind, and the default mode: a quantitative meta-analysis. J Cogn Neurosci. 2009;21: 489–510. doi: 10.1162/jocn.2008.21029.

17. Szpunar KK, Watson JM, McDermott KB. Neural substrates of envisioning the future. Proc Natl Acad Sci. 2007; 104: 642–647. doi: 10.1073/pnas.0610082104.

18. Svoboda E, McKinnon MC, Levine B The functional neuroanatomy of autobiographical memory: A meta-analysis. Neuropsychologia. 2006;44: 2189–2208. doi: 10.1016/j.neuropsychologia.2006.05.023.

19. Rugg MD, Vilberg KL. Brain networks underlying episodic memory retrieval. Curr Opin Neurobiol. 2013;23: 255–260. doi: 10.1016/j.conb.2012.11.005.

20. Cabeza R, St Jacques P Functional neuroimaging of autobiographical memory. Trends Cogn Sci. 2007;11: 219–227. doi: 10.1016/j.tics.2007.02.005.

21. Raichle ME, MacLeod AM, Snyder AZ, Powers WJ, Gusnard DA, Shulman GL. A default mode of brain function. Proc Natl Acad Sci. 2001; 98: 676–682. doi: 10.1073/pnas.98.2.676.

22. Buckner R, Andrews-Hanna JR, Schacter DL. The Brain9s Default Network: Anatomy, Function, and Relevance to Disease. Ann N Y Acad Sci. 2008; 1–38.

23. Kriegeskorte N, Goebel R, Bandettini P Information-based functional brain mapping. Proc Natl Acad Sci U S A. 2006; 103: 3863–3868. doi: 10.1073/pnas.0600244103.

24. Kriegeskorte N, Mur M, Bandettini P, Representational Similarity Analysis – Connecting the Branches of Systems Neuroscience. Front Syst Neurosci. 2008;2. doi: 10.3389/neuro.06.004.2008.

25. Maguire EA. Neuroimaging studies of autobiographical event memory. Philos Trans R Soc Lond Ser B. 2001;356: 1441–1451. doi: 10.1098/rstb.2001.0944.

26. Wing EA, Ritchey M, Cabeza R, Reinstatement of Individual Past Events Revealed by the Similarity of Distributed Activation Patterns during Encoding and Retrieval. J Cogn Neurosci. 2014; 1–13. doi: 10.1162/jocn_a_00740.

27. Dikker S, Silbert LJ, Hasson U, Zevin JD. On the Same Wavelength: Predictable Language Enhances Speaker–Listener Brain-to-Brain Synchrony in Posterior Superior Temporal Gyrus. J Neurosci. 2014;34: 6267–6272. doi: 10.1523/JNEUROSCI.3796-13.2014.

28. Hassabis D, Kumaran D, Maguire EA. Using Imagination to Understand the Neural Basis of Episodic Memory. J Neurosci. 2007;27: 14365–14374. doi: 10.1523/JNEUROSCI.4549-07.2007.

29. Buckner RL, Carroll DC. Self-projection and the brain. Trends Cogn Sci. 2007;11: 49–57. doi: 10.1016/j.tics.2006.11.004.

30. Honey CJ, Thompson CR, Lerner Y, Hasson U. Not lost in translation: neural responses shared across languages. J Neurosci Off J Soc Neurosci. 2012;32: 15277–15283. doi: 10.1523/JNEUROSCI.1800-12.2012.

31. Regev M, Honey CJ, Simony E, Hasson U. Selective and Invariant Neural Responses to Spoken and Written Narratives. J Neurosci. 2013;33: 15978–15988. doi: 10.1523/JNEUROSCI.1580-13.2013.

32. Binder JR, Desai RH, Graves WW, Conant LL. Where is the semantic system? A critical review and meta-analysis of 120 functional neuroimaging studies. Cereb Cortex N Y N 1991. 2009;19: 2767–2796. doi: 10.1093/cercor/bhp055.

33. Hassabis D, Kumaran D, Vann SD, Maguire EA. Patients with hippocampal amnesia cannot imagine new experiences. Proc Natl Acad Sci U S A. 2007; 104: 1726–1731. doi: 10.1073/pnas.0610561104.

34. Romero K, Moscovitch M. Episodic memory and event construction in aging and amnesia. J Mem Lang. 2012;67: 270–284. doi: 10.1016/j.jml.2012.05.002.

35. Ranganath C, Ritchey M. Two cortical systems for memory-guided behaviour. Nat Rev Neurosci. 2012;13: 713–726. doi: 10.1038/nrn3338.

36. Zwaan RA, Radvansky GA. Situation models in language comprehension and memory. Psychol Bull. 1998;123: 162–185.

37. Zacks JM, Speer NK, Swallow KM, Braver TS, Reynolds JR. Event Perception: A Mind/Brain Perspective. Psychol Bull. 2007;133: 273–293. doi: 10.1037/0033-2909.133.2.273.

38. Baldassano C, Chen J, Zadbood A, Pillow JW, Hasson U, Norman KA. Discovering event structure in continuous narrative perception and memory. bioRxiv. 2016; 081018. doi: 10.1101/081018.

39. Cichy RM, Heinzle J, Haynes J-D. Imagery and perception share cortical representations of content and location. Cereb Cortex N Y N 1991. 2012;22 372–380. doi: 10.1093/cercor/bhr106.

40. Johnson MR, Johnson MK. Decoding individual natural scene representations during perception and imagery. Front Hum Neurosci. 2014;8: 59. doi: 10.3389/fnhum.2014.00059.

41. Stokes M, Thompson R, Cusack R, Duncan J. Top-down activation of shape-specific population codes in visual cortex during mental imagery. J Neurosci Off J Soc Neurosci. 2009;29: 1565–1572. doi: 10.1523/JNEUROSCI.4657-08.2009.

42. Vetter P, Smith FW, Muckli L. Decoding Sound and Imagery Content in Early Visual Cortex. Curr Biol. 2014;24: 1256–1262. doi: 10.1016/j.cub.2014.04.020.

43. Bartlett FC, Burt C. Remembering: A Study in Experimental and Social Psychology. Br J Educ Psychol. 1933;3: 187–192. doi: 10.1111/j.2044-8279.1933.tb02913.x.

44. John D. Bransford MKJ. “Contextual Prerequisites for Understanding: Some Investigations of Comprehension and Recall.” J Verbal Learn Verbal Behav. 1972;11: 717–726. doi: 10.1016/S0022-5371(72)80006-9.

45. Alba JW, Hasher L. Is memory schematic? Psychol Bull. 1983;93: 203–231. doi: 10.1037/0033-2909.93.2.203.

46. Clark HH. Context and Common Ground. In: Brown K, editor. Encyclopedia of Language and Linguistics. Elsevier; 2006. pp. 105–108.

47. Clark HH, Krych MA. Speaking while Monitoring Addressees for Understanding. J Mem Lang. 2004;50: 62–81. doi: 10.1016/j.jml.2003.08.004.

48. Pickering MJ, Garrod S. Toward a mechanistic psychology of dialogue. Behav Brain Sci. 2004;27: 169-190-226.

49. Binder JR, Desai RH. The neurobiology of semantic memory. Trends Cogn Sci. 2011;15: 527–536. doi: 10.1016/j.tics.2011.10.001.

50. Margulies DS, Ghosh SS, Goulas A, Falkiewicz M, Huntenburg JM, Langs G, et al. Situating the default-mode network along a principal gradient of macroscale cortical organization. Proc Natl Acad Sci. 2016; 201608282. doi: 10.1073/pnas.1608282113.

51. Hasson U, Chen J, Honey CJ. Hierarchical process memory: memory as an integral component of information processing. Trends Cogn Sci. 2015;19: 304–313. doi: 10.1016/j.tics.2015.04.006.

52. Benjamini Y, Hochberg Y. Controlling the False Discovery Rate: A Practical and Powerful Approach to Multiple Testing. J R Stat Soc Ser B Methodol. 1995;57: 289–300.

53. Hasson U, Nir Y, Levy I, Fuhrmann G, Malach R. Intersubject Synchronization of Cortical Activity During Natural Vision. Science. 2004;303: 1634–1640. doi: 10.1126/science.1089506.

54. Shirer WR, Ryali S, Rykhlevskaia E, Menon V, Greicius MD. Decoding Subject-Driven Cognitive States with Whole-Brain Connectivity Patterns. Cereb Cortex. 2011;bhr099. doi: 10.1093/cercor/bhr099.

